# Detergent selection as a determinant of protein melting behavior and drug-target interaction-calling in thermal stability proteomics

**DOI:** 10.64898/2026.05.28.728510

**Authors:** Catherine Sniezek, Veronika A. Glukhova, Christian Schmitz, Katarina Vlajic, Devin K. Schweppe

**Affiliations:** Department of Genome Sciences, University of Washington, Seattle, WA 98105, USA; Brotman Baty Institute for Precision Medicine, Seattle, WA 98105, USA; Institute for Stem Cell & Regenerative Medicine, University of Washington, Seattle, WA 98105, USA

## Abstract

Thermal proteome profiling (TPP) and proteome integral solubility alteration (PISA) assays measure drug–target interactions by monitoring protein thermal stability across the proteome. While detergents are routinely used in lysate-based thermal profiling, the field lacks consensus on whether detergents should be present during the melting step or only added afterward as an extraction buffer, and whether detergent identity matters for this choice. Here, we evaluate how commonly used detergents and the timing of their use in thermal stability workflows affect proteome-wide thermal stability and PISA hit calling in TF-1 lysates. We find that NP-40 and DDM produce highly correlated melting profiles when used exclusively as post-melt extraction buffers, but diverge substantially when present during the melting step. DDM in particular prevents the thermally-induced loss in solubility of large classes of proteins, such as cell surface proteins, and these effects propagate directly into PISA hit calling. Performing the PISA melt in DDM versus NP-40 results in the gain and loss of distinct drug–target interactions for both the PAK4 inhibitor PF-3758309 and the PLK1 inhibitor volasertib. Notably, DDM enables detection of a volasertib–TMEM97 interaction that was previously not detected in NP-40. However, we also find that the stabilization effects of DDM mask the identification of some known PISA hits for these drugs. We further introduce a four-parameter logistic model of protein melting to aid in modeling of these findings and a linear regression framework for PISA hit calling that outperforms pairwise t-tests in low-replicate settings. Together, these results establish detergent selection as a tunable experimental variable in thermal profiling and suggest that some drug–target engagements previously attributed exclusively to intact-cell context may be recoverable in lysates with appropriate buffer conditions.

## Introduction

Thermal proteome profiling (TPP)^1^ and the condensed, high-throughput proteome integral solubility alteration (PISA)^2^ assays have emerged as tools for probing a proteome’s response to stimuli. Both assays work on the principle that proteins have a characteristic melting point (Tm) at which they denature and unfold, potentially leading to aggregation and precipitation. Comparison of proteome-wide melting behaviors with and without ligands or drugs then enables detection of ligand- and drug-engagement across the proteome ^1^. As such, the ability to sensitively measure changes in thermal stability is a critical aspect of thermal profiling.

Thermal profiling experiments vary in the type and order of detergent use, specifically in regards to the melting step. Thermal profiling on intact cells necessitates melting in gentle buffer conditions, typically PBS with minimal additives to preserve cell membrane integrity and ensure physiological relevance of drug-target engagement. On the other hand, thermal profiling in cell lysates permits flexibility in application of additives such as detergent. However, detergents have not been applied consistently across thermal profiling experiments. Some protocols mechanically lyse cells in PBS, perform the melt in PBS, and then treat with detergent after the melting step to improve protein recovery^3–5^. Other protocols lyse cells in the presence of detergent, therefore necessitating that the melting step occurs in detergent^6–8^. Still other protocols include both methods for lysate prep steps in their methods^9^. Given the inconsistency in the literature for the order of detergent application for thermal profiling on cell lysates, we reasoned that a systematic evaluation of detergent use was needed, specifically, an investigation of whether and how detergents alter thermal melting of proteins.

A successful lysis buffer component for thermal profiling experiments can be evaluated by a few key metrics. First, the detergent must be either mass-spec compatible or easily removed by sample cleanup. Second, the detergent must efficiently extract and solubilize proteins for downstream analysis. Finally, for thermal profiling particularly, a good detergent will solubilize non-denatured proteins but will not interfere with the aggregation and precipitation process of heat-denatured proteins. The nonionic detergent NP-40 has been well-characterized in both cell-based and lysate-based thermal profiling^6^ and PISA assays^3^. More recently, the detergent n-dodecyl-beta-maltoside (DDM) has been demonstrated as a viable option as a post-melt additive for high-throughput thermal stability assays^5^, and DDM’s compatibility with mass spectrometry has permitted its recent increased use in proteomics assays across the board^10,11^.

Here, we characterize the efficacy of DDM and NP-40 as lysis buffer components in thermal profiling experiments. We find that NP-40 and DDM produce comparable proteome-wide melting profiles when used only as post-melt extraction buffers, but diverge substantially when present during the melting step. These detergent-dependent differences propagate directly into PISA hit calling, showing that drug–target interactions can be gained or lost depending on lysis buffer choice alone. Together, these results suggest that detergent selection should be treated as a tunable experimental variable in thermal profiling. By tuning buffer conditions and improving statistical sensitivity, we demonstrate in the case of CHEK1 and bromodomain proteins BRD2/3 that we can recover drug–target engagements in lysates that were previously thought to require an intact cellular context.

## Methods

### Cell culture and lysate preparation

TF-1 erythroleukemia cells (ATCC CRL-2003) were grown under standard tissue culture conditions. Media used was RPMI (brand) with 2 ng/mL GM-CSF (brand), 10% FBS (brand), and pen/strep (specifics, brand). Cells were maintained at 1-8 x 10^5^ cells/mL, and were harvested by centrifugation, washed twice with PBS, and stored at −80°C until use. For each condition, frozen pellets containing 1 × 10⁷ cells were resuspended in ice-cold PBS supplemented with protease inhibitor (Roche cOmplete Mini, EDTA-free) and adjusted with an equal volume of 2× lysis solution to a final volume of 50 µL. 2× lysis solutions were prepared in PBS with protease inhibitor and yielded the following final conditions: 0.5% or 1% NP-40 (Thermo Scientific NP-40 Surfact-Amps, Cat. 85124), 0.5% DDM (Thermo Scientific n-dodecyl-β-D-maltoside, 99%, Cat. 329370010), 1% SDS (Sigma, Cat. 851922), 0.5% NP-40 + 0.5% DDM, 0.5% LMNG (Thermo Scientific lauryl maltose neopentyl glycol, Cat. A50940), and 0.5% NM (Thermo Scientific n-nonyl-β-maltoside, Cat. A65513). Lysates were incubated on ice for 15 min, then subjected to heat treatment at either 37°C or 70°C for 3 min, immediately transferred back to ice, and centrifuged at 20,000 × g for 2 h at 4°C. The soluble supernatant was separated from the aggregated pellet by pipetting.

### SDS-PAGE and western blots

Cell lysates were combined with 4× loading buffer (Novex LDS Sample Buffer, Cat. NP0008) to a final volume of 25 µL, heated at 95°C for 5 min, cooled, and resolved on 4–12% Bis-Tris gels (Novex Bolt 4–12% Bis-Tris Plus WedgeWell, Cat. NW04120BOX) in MOPS running buffer (NuPAGE MOPS, 20x, Cat. NP0001) at 120 V for 50 min. Separated proteins were transferred and immobilized onto PVDF membranes (Novex, Cat. IB24001) using the iBlot 2 Gel Transfer Device (ThermoFisher Scientific, Cat: IB21001) according to manufacturer’s instructions.

All subsequent wash steps were performed in TBS-T buffer: 1x TBS (ThermoFisher Scientific TRIS-buffered saline, 10x, pH 7.4, Cat: J60764.K2) and 1% v/v Tween-20 (Millipore Sigma Tween 20, 40% stock solution, Cat: P9416). Membrane was blocked in solution of 5% nonfat dry milk in TBS-T for 2 hours at RT. Primary antibodies against TNFRSF1A (Cell Signaling Technology, Cat: 3736T), TrkA (Invitrogen, Cat. MA5-32123), AurkB (CST, Cat. 28711), beta-Actin (CST, Cat. 4970), Na K ATPase (Bethyl Laboratories, Cat. A500-031A-T), TMEM97 (CST, Cat. 62790), was diluted 1:1000 in 5% nonfat dry milk in TBS-T and incubated at 4°C overnight with shaking. After washing 3x in TBS-T, the membrane was incubated with anti-rabbit secondary antibody (CST, Cat. 7074P2) at 4°C for 1 hr with shaking. Membrane was washed 3x with TBS-T prior to imaging on Bio-Rad ChemiDoc imaging system.

### Isothermal melts

Frozen TF-1 cells were thawed in a minimal amount of PBS. For the PBS melt, cells were brought to 5 million cells/mL with PBS plus protease inhibitors (Roche cOmplete, EDTA-free mini tablets). For the detergent melt, cells were split into two aliquots and brought to 5 million cells/mL in either 0.5% NP-40 or 0.5% DDM in PBS plus protease inhibitor. Resulting crude lysates were aliquoted at 100 µL in triplicate into PCR strips. Lysates were heated to 37, 50, or 70°C in a PCR thermocycler held at 25°C for 3 minutes, followed by 3 minutes of heating, then cooled to 25°C for 3 minutes. After heating, PBS melt samples were treated with a 2x lysis buffer (final composition 0.5% NP-40 or 0.5% DDM in PBS plus protease inhibitor). All samples were then centrifuged at 20,000 xg for 2 hours, and supernatants were collected for downstream analysis.

### Drug treatment and PISA assay

Frozen TF-1 cells were thawed in a minimal amount of PBS and aliquoted into two portions. Cells were resuspended in either 0.5% NP-40 or 0.5% DDM in PBS plus protease inhibitor (Roche cOmplete, EDTA-free mini tablets) and mechanically lysed with pipetting. Lysate was cleared by centrifugation at 20,000 xg for 30 minutes. Cleared lysate was divided into 6 aliquots for the six conditions: NP-40 lysis plus DMSO, PF-3758309 (10 µM, Med Chem Express HY-13007) or volasertib (10 µM, Med Chem Express HY-12137); DDM lysis plus DMSO, PF3758309 (10 µM) or volasertib (10 µM). Treated lysates were allowed to incubate for 15 minutes. Meanwhile, 15 µL of treated lysate was aliquoted into 10 wells of a PCR plate such that each treatment received, in triplicate, temperatures ranging from 48-58°C in a thermocycler. Thermocycler was set to hold at 25°C for 3 minutes, followed by 3 minutes of gradient heating from 48-58°C, then 3 minutes at 25°C. Per treatment condition, the 10 aliquoted temperature points were pooled and centrifuged at 20,000 xg for 2 hours. Resulting supernatants were collected for downstream analysis.

### Sample workup for TMT proteomics

All samples were worked up for standard TMT quantitative proteomics. Protein content of cleared supernatants was quantified using BCA (Pierce). 20 ug of protein was treated with a reducing buffer (final composition 0.5% SDS, 5 mM TCEP in EPPS, pH 8.5) for 10 minutes. Samples were then treated with 10 mM IAA for 20 minutes, then 10 mM DTT for 10 minutes. Resulting reduced and alkylated samples were cleaned by SP3^12^. A 1:1 mixture of Mag4 and Mag6 SP3 beads (Cytiva) was prepared by cleaning twice with water. Beads were added to sample at a 10:1 mass ratio of beads:protein, then to precipitate proteins 2x sample volume of 100% ethanol was added so that final ethanol content was 66%. Precipitating proteins were vortexed for 30 minutes, and then cleaned by 3 subsequent washes of 80% ethanol. Finally, beads were resuspended in a digestion solution containing 1:100 enzyme:protein of both trypsin (Promega, sequencing grade) and LysC (Wako) in EPPS 8.5. Digestion occurred overnight at 37°C.

Resulting cleaned peptides were labelled with TMTpro (Thermo Fisher) at a ratio of 2.5:1 TMT:peptide in the presence of 35–40% acetonitrile for 1 hour. A small amount (approximately 10%) of labelled peptides was pooled and analyzed by mass spectrometry for physical normalization. Samples were pooled such that equal amounts of labelled peptides from each channel, as determined by mass spectrometry, were combined, and cleaned by Sep-Pak (Waters Oasis Sep-Pak tC18. Cat. WAT054960)). The resulting pooled and cleaned sample was fractionated by reverse-phase HPLC (RP-HPLC)^13^. Mobile phase A was 5% acetonitrile, 10 mM ammonium bicarbonate pH 8, mobile phase B was 90% acetonitrile, 10 mM ammonium bicarbonate pH 8, with a C18 column (Zorbax 300Extend-C18, 4.6×250mm, 5-micron) and a flow rate of 0.6 mL/min. Gradient started at 100% A, then 1-44% B from 2-60 minutes, followed by a 5 minute wash at 99% B and a 10 minute equilibration at 99% A. 96 fractions were collected from 5-75 minutes, which were pooled into 24 fractions, and cleaned by STAGE-tip^14^ before mass spectrometry analysis.

### Mass spectrometric methods

All samples were analyzed with an Orbitrap Eclipse Tribrid (Thermo Fisher) with FAIMS installed and with an Easy nLC (Thermo Fisher) equipped with a C18 analytical column (15 cm Odyssey, ionOpticks). Peptides were resuspended in 5% acetonitrile, 2% formic acid and separated with a 90 minute chromatography method which started at 96% mobile phase A (5% acetonitrile, 0.125% formic acid), 4% mobile phase B (95% acetonitrile, 0.125% formic acid) followed by a 75 minute gradient to 28% B and a subsequent wash for 10 minutes at 100% B. Peptides were analyzed and TMT quantified with a real-time search-enabled (RTS)^15–17^, single-precursor selection (SPS) MS3 approach. Precursors were detected in MS1 (Orbitrap resolution 120,000, 200% normalized AGC, maximum injection time 50 ms, FAIMS CVs -40, -60, -80V; isolation range 400-2000 m/z). Precursors were collected for MS2 fragmentation and analysis (quadrupole isolation, ion trap CID 35%, 200% normalized AGC, 75 ms inject time). Dynamic exclusion was enabled with a 90s delay, a 10 ppm window, and intensity filter greater than 5,000. Real-time search was applied to MS2 spectra against the human protein database, static carbamidomethyl and TMTpro modifications, and variable oxidized methionines. RTS scoring thresholds were 2 Xcorr, 0.1 dCn, 10 ppm precursor ppm, and charge state of 2. Synchronous precursor selection was used with n=10 fragments selected for MS3 fragmentation and analysis for TMTpro label quantification (first mass 110 m/z, Orbitrap resolution 50,000, 400% normalized AGC, HCD 45%, 120 ms max inject time). Spectra were searched offline with the Comet search algorithm ^18^ and filtered to a 1% protein and peptide false discovery rate using linear discriminator analysis and rules of protein parsimony^16^. Proteins were quantified using relative quantification of SPS-MS^3^ reporter ions and normalized with an equalized-medians approach. Raw files and searches available on ProteomeXchange with identifier PXD078871.

### Post-hoc data analysis

Protein melting simulations were performed and visualized with R (version 4.5.1). Protein solubility for each integer temperature point from 35-75°C was modeled using the following equation, which is an adaptation of the logistic function:

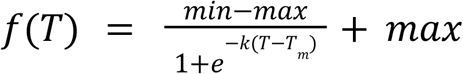

Where *T* is temperature, and the four constants are defined as: *min* is minimum plateau of protein solubility, *max* is maximum plateau of protein solubility, *k* is a slope constant, and *T_m_* is the midpoint of the function, here defined as the melting point. Except where defined, default constants were set to min = 0, max = 1, k = 0.5, and Tm = 50°C. To determine default constant parameters, the above equation was fit to a portion of the human Meltome dataset^9^ with a nonlinear least squares approach. Initial fit conditions were *min* = 0.01, *max* = 1, *k* = 0.5, and *Tm* = 52°C and boundary conditions were set at min > 0.001. This curve fit approach was also used to model detergent effects on thermal profiling from a publicly available TPP set^6^.

Protein quantitation matrices were manipulated and visualized using R (version 4.5.1) and normalized with COMBAT batch correction^19^. For isothermal melts, the B1 slope coefficient was calculated using the following equation:

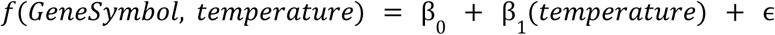

Proteins ranked by melting slope behavior either by difference of detergent order (detergent melt - PBS melt) or by different of detergent type (DDM - NP-40) and were evaluated with fgsea^20^ with minSize set to 5, maxSize set to 500. The pfam protein domain dataset^21^ was obtained by downloading all human enrichment terms from the String database^22^ (enrichment terms version 12.0) and filtering for the Pfam category. Jensen’s COMPARTMENTS enrichment dataset^23^ was downloaded directly from the COMPARTMENTS website (https://compartments.jensenlab.org/Downloads, November 7, 2025).

For PISA data, p-values of treatment versus DMSO controls were determined in three different ways for comparisons as specified: Welch’s t-test, Limma^24^, or a linear model approach with the following equation:

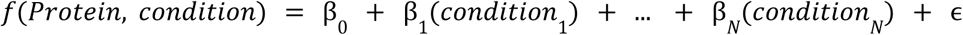

Where each condition is a different detergent by drug treatment. For NP-40 analysis, condition NP-40-DMSO was set as reference. For DDM analysis, condition DDM-DMSO was set as reference. All P-values were corrected by the Benjamini-Hochberg approach^25^ before reporting. Fold-change cutoffs were set at FC > 1.25 or < 1/1.25.

For comparisons with Van Vranken, 2024 results, supplemental data was downloaded and TMT plexes containing live-cell or cell extract conditions for volasertib or PF-3758309 treatment were identified. In these sets, significance of thermal shifts was determined by examining the distribution of all measured thermal shifts and defining standard deviations from this distribution. For this current work, proteins were defined as hits in the Van Vranken dataset if they had |SD| > 3.5 and fold-change cutoff of FC > 1.25 or < 1/1.25.

Code for the analyses performed here is available at https://github.com/SchweppeLab/DDM_PISA_paper. Claude Sonnet 4.6 was used to aid in bug fixes, data matrix manipulations, and data visualizations.

## Results & Discussion

### Determinants of thermal profiling

Identification of drug–target interactions by thermal profiling rests on the premise that proteins possess a mutable thermal stability that can be shifted by ligand binding. A less appreciated aspect of these experiments is the extent to which the assay buffer itself sets the baseline thermal state of the proteome, and therefore gates which thermal stability shifts are measurable in the first place. Beyond the inherent protein sequence and structure effects, protein thermal stability is a relative property, governed in part by environmental factors such as pH, ionic strength, and co-solutes. Because temperature-induced protein aggregation is thought to be driven predominantly by hydrophobic interactions^26,27^, detergents, which reshape the hydrophobic environment of the lysate, should be expected to alter aggregation behavior^28^.

Protein melting is a thermodynamic process which can be modelled from first principles as a logistic function^1^ (**Fig 1B**). Current methods of thermal profiling data analysis use a three-parameter function to model protein solubility as a function of temperature^1,8,9^. However, a four parameter logistic function can also be used, with the four fit parameters as descriptions of melting behavior: minimum protein quantitation *min*, maximum protein amount *max*, a slope parameter *k*, and melting point *Tm* modelled directly from the fit curve (**Fig 1C**). Whereas previous methods define melting slope and melting points post-hoc with first and second derivatives, respectively^1^, this new method has the benefit of modeling these variables directly. Fitting a melting curve across thousands of proteins in a publicly-available thermal profiling dataset^9^ (**Fig S1A**) resulted in curves with excellent agreement with the source data (median Rsqd = 0.99), with easily identifiable proteins with good melting behavior (**Fig S1B**), high minimum protein stability (**Fig S1C**), or shallow melting slopes (**Fig S1D**). Using the median values of this dataset as a starting point, we modeled how changing each of the four parameters altered the shape of the melting curve and resulting fold change of a five-degree melting point shift (**Fig 1D-G**). With this approach, we find several conditions are needed for optimal fold-changes in an integrated thermal profiling experiment such as PISA. First, the protein must lose its solubility to completion at high temperature (**Fig 1E**). Second, the protein must melt and aggregate rapidly around its melting point (**Fig 1F**), and finally the initial melting point of the protein must fall within the melting temperature window being used, or ideally before the window in the case of a positive expected thermal shift (**Fig 1G**). Loss of any of these conditions should lead to smaller quantitative effect sizes due to ligand binding and therefore have the potential to cause false negatives in a PISA assay.

**Figure 1.**
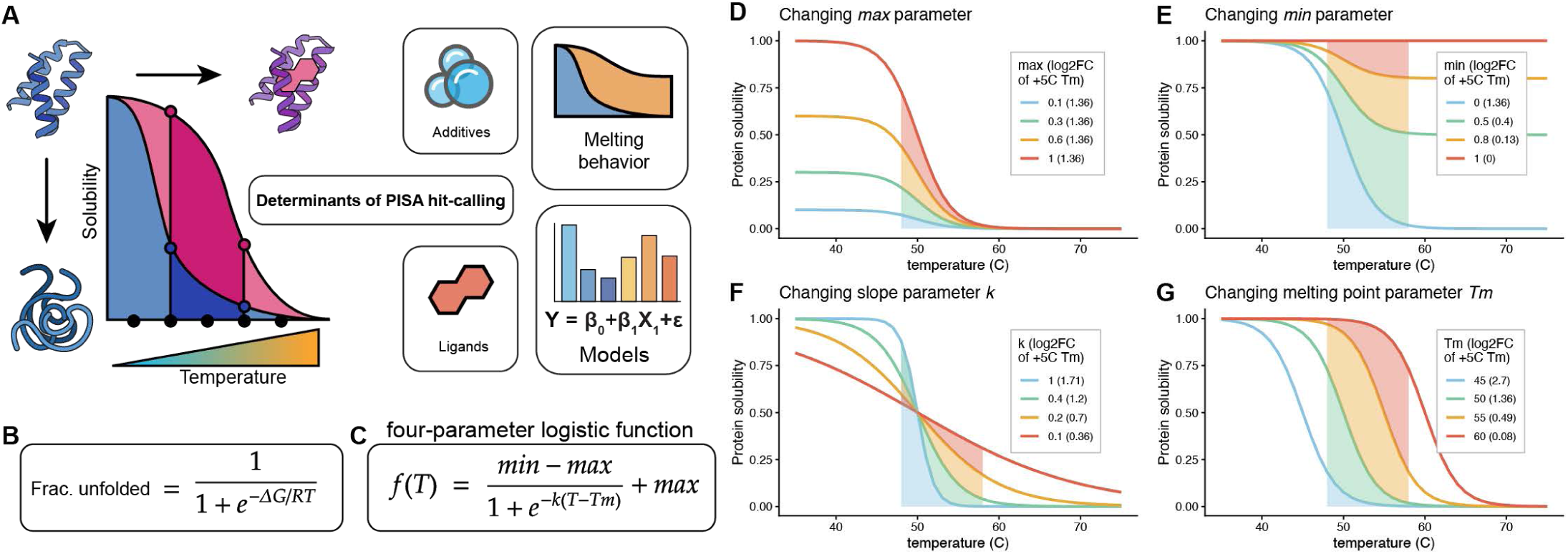
Determinants of hit-calling in thermal profiling assays. (A) Investigation of the determinants of thermal stability proteomics hit calling based on combinatorial analysis of detergent addition and ligand engagement. (B) Theoretical melting curve as a function of temperature derived from protein unfolding kinetics. (C) Four-parameter logistic function modelling protein solubility as a function of temperature, with min, max, and slope k constants. (D-G) Alterations to theoretical melting curves by changing maximum protein amount (D), minimum protein amount (E), shape of melting profile (F) or intrinsic melting point (G). Default parameters derived from median values for four-point curve fit across a publicly-available TPP dataset^9^ (**Fig S1**) (max = 1, min = 0, k = 0.5, Tm = 50). The shaded area indicates a 48-58°C PISA integration window. Predicted log2FC calculated by adding +5°C to melting point Tm and integrating resulting curves from 48-58°C.

### Detergent use impacts lysate proteome thermal stability

Given that ideal protein melting behavior is a condition for optimal thermal profiling, and given that detergents are expected to alter the thermal stability of proteins, we systematically evaluated the impact of detergent type and order of use on the thermal stability of the lysate proteome. Frozen-thawed crude TF-1 lysate prepared in PBS was subjected to isothermal melts at 37, 50, and 70°C either without detergent or in the presence of 0.5% NP-40 or 0.5% DDM. These concentrations were chosen as they are routinely used in lysate-based TPP/PISA workflows for NP-40^3,6,8^ and elicited differential stabilization of membrane proteins TNFR1 and NTRK1 (**Fig. S2C**). To isolate the direct effect of detergent on melting and to mirror existing lysate-based thermal profiling protocols ^3–7,9^, all samples received 0.5% NP-40 or 0.5% DDM after the melt as an extraction buffer (**Fig. 2A**).

**Figure 2.**
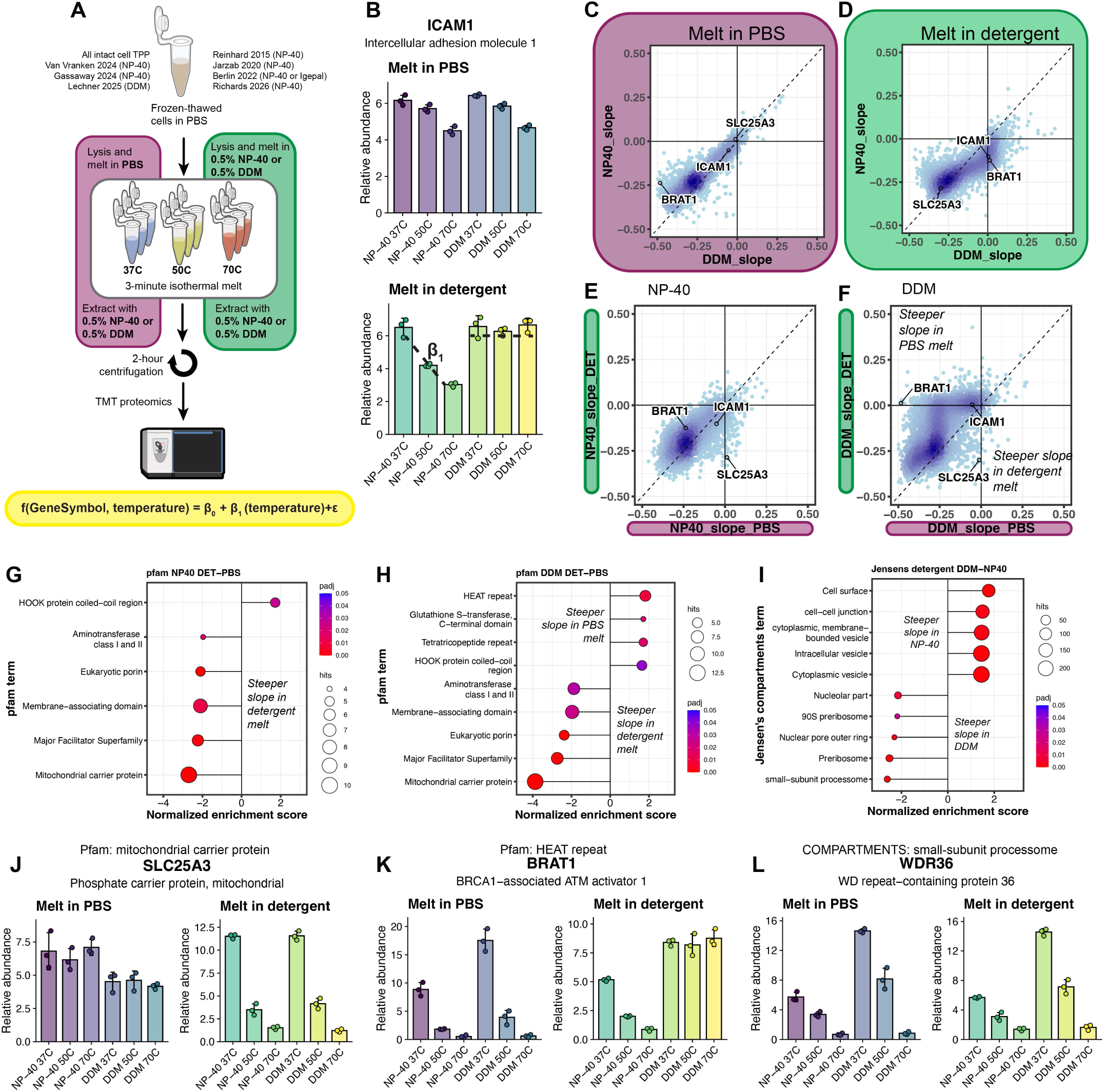
Melting in detergent alters thermal profiles of lysate proteome. (A) Overview of melt experiments which compare melting in the presence of detergent (green) with melting in the presence of PBS, followed by detergent extraction (pink). (B) Melting profiles of cell-surface protein ICAM1 when melted in PBS and when melted in detergent, with melting slope β1 highlighted. (C) β1 melting slopes for PBS melt with each extraction detergent directly compared. (D) β1 melting slopes for detergent melts with each detergent compared. (E) NP-40 β1 melting slopes for detergent melt or PBS melt. (F) DDM β1 melting slopes for detergent melt or PBS melt. (G-H) Proteins ranked by different of β1 slopes (detergent melt - PBS melt) for NP-40 (G) or DDM (H) and analyzed with fgsea^20^ with pfam protein domain database^21^. Terms were ranked by adjusted P-value and up to 5 terms with p.adj < 0.05 were selected for each positive or negative NES direction. Negative enrichment scores represent steeper melting profile when melted in detergent. (I) Proteins ranked by difference of β1 melting slopes in the presence of detergent (DDM melt - NP-40 melt) and analyzed with fgsea with Jensen’s compartments database^23^.

Using the isothermal melt measurements, relative temperature effects were estimated for each quantified protein with a linear model, using the temperature coefficient (β1) as a proxy for melting behavior (**Fig. 2B**). Negative β1 values indicated loss of solubility with increasing temperature, while slopes near zero indicated little to no melting. When detergent was used only as a post-melt extraction buffer, DDM- and NP-40-treated samples produced highly similar β1 distributions (**Fig. 2C**; Pearson *r* = 0.92). This was in agreement with the finding that NP-40 use post-melt, compared to PBS alone, had little effect on the melting behavior of proteins^6^. In contrast, when lysate was treated with detergent before melting, DDM and NP-40 treated samples diverged substantially (**Fig. 2D**), with DDM producing a bimodal distribution of melting coefficients that was absent under NP-40 (**Fig. S3F**). Comparing each detergent against its PBS-only control made this difference asymmetry explicit: NP-40 yielded similar melting behavior whether used only as an extraction buffer or as a melt buffer (**Fig. 2E**; Pearson *r* = 0.71), whereas DDM produced markedly different profiles when present during the melt itself (**Fig. 2F**; Pearson *r* = 0.64). Many proteins, including ICAM1, melted in NP-40 but not in DDM (**Fig. 2B, Fig S3G-I**). Conversely, a substantial set of proteins, including TMEM97, exhibited clear melting behavior only when detergent was present (**Fig S3G, Fig S4A, Fig. 5F**). Our data confirm the expectation that detergents which reshape the hydrophobic environment of lysates alter aggregation behavior, and show that melting in detergent, DDM in particular, changes the aggregation trajectories of a substantial fraction of the proteome.

To identify which classes of proteins were most sensitive to detergent choice during the melt step, we ranked proteins based on the difference in temperature slope (detergent-melt β1 − PBS-melt β1); negative values indicated steeper slopes, and therefore more temperature-induced aggregation when melted in detergent versus in PBS. We then used this ranked list for gene-set enrichment analysis with fgsea^20^ against the Pfam domain gene set^21^ (**Fig. 2G-H**). Both detergents enhanced temperature-induced aggregation of mitochondrial carrier proteins, membrane-associating domains, and porins, with DDM showing a stronger effect than NP-40. This is consistent with the reported behavior of these detergents toward integral membrane proteins: detergent micelles solubilize membrane proteins and in some cases preserve native protein conformations.^28–30^. DDM additionally reduced aggregation for several domain classes that NP-40 did not systematically affect, including HEAT and tetratricopeptide repeats, elongated α-helical scaffolds whose extensive hydrophobic surfaces are normally buried within large multiprotein complexes, and which may therefore be better shielded in the presence of DDM (**Fig. 2H**).

Further, we explored classes of proteins with a change in thermal stability that was dependent on the detergent used during the melt (DDM-melt β1 − NP-40-melt β1). In general, DDM produced steeper melting profiles for ribosomal proteins and nuclear pores, while NP-40 shows a strong effect on membrane-bound proteins (**Fig. 2I**), principally driven by proteins which exhibited more melting in the presence of NP-40 compared to PBS and considerably less melting in DDM compared to PBS (**Fig. S2)**. This finding was consistent with a model that DDM improved extraction efficiency of some proteins such as nuclear proteins, yet the stabilization effect on membrane proteins prevented the thermally-induced aggregation of such proteins. Together, these data show that while NP-40 and DDM produce comparable profiles when used strictly as extraction buffers, they exert markedly different effects on proteome thermal stability when present during the melt.

### Detergent use impacts PISA hit identification

Based on the different melting profiles of proteins melted in DDM compared to NP-40, we performed PISA assays in the presence of the two detergents to identify how they affect hit-calling for protein-ligand interactions. Frozen-thawed TF-1 cells were resuspended with either 0.5% DDM or 0.5% NP-40 and the resulting lysates were treated with DMSO, Volasertib (PLK1/BRD inhibitor), or PF-3758309 (PAK4 inhibitor, hereafter ‘PF-37’) (**Fig. 3A**). Following drug treatment, lysates were aliquoted into 10 temperature points ranging from 48-58°C, then pooled for physical integration and centrifuged to remove insoluble, aggregated proteins. Volasertib and PF-37 were selected for their known interactions with proteins which we found to have differential melting behavior in the isothermal melts. For example, PF-37 targets DDM-stabilized STK3 (**Fig 3B**) and DDM-destabilized CHEK1 (**Fig 3C**), and volasertib targets DDM-stabilized BRD3 (**Fig 4H**). Further, volasertib has historically only produced PISA hits in an in-cell context, while PF-37 produces hits in both lysate and in-cell PISA contexts^3^. This compound pair therefore allowed us to probe detergent effects on PISA hit calling from two complementary angles: compounds with cell-context-dependent and cell-context-independent engagement.

**Figure 3.**
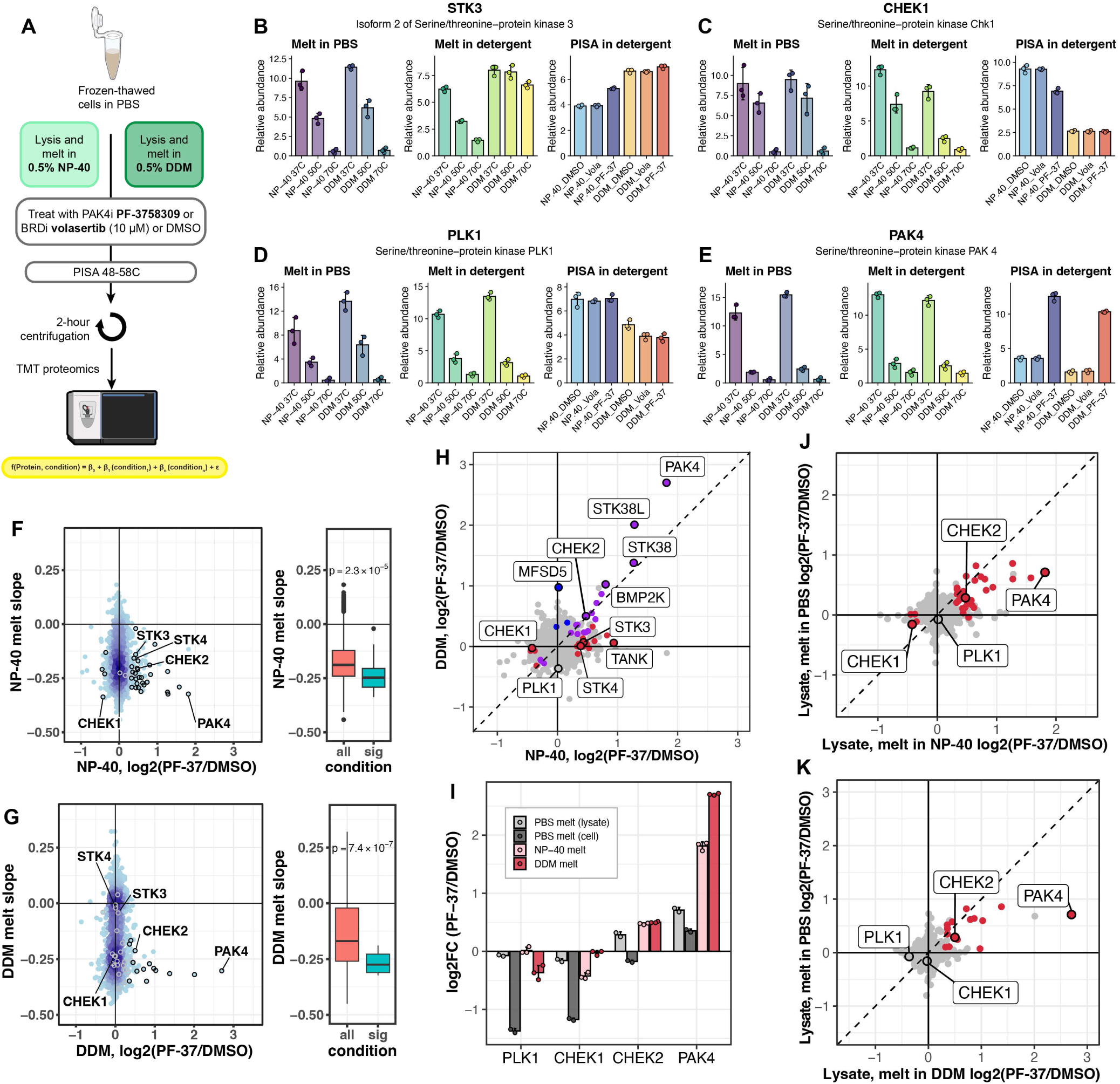
Altered melting profiles of detergent melts impact PISA drug-target identifications. (A) Overview of experiment, samples were lysed in either NP-40 or DDM and treated with either the PAK4i PF-3758309 (‘PF-37’) or the PLKi volasertib. (B-E) Selected proteins with melting profiles and PISA quantitative results, demonstrating DDM-stabilization and loss of PISA hit for STK3 (B), DDM-destabilization and loss of hit for CHEK1 (C), known PF-37 hit PLK1 (D), and largest fold-change across both conditions PAK4 (E). (F-G) Temperature β1 melting slopes from Figure 1 plotted against log2(treatment/DMSO) thermal shifts from PISA assay. Black circles indicate significant PISA hit for that condition (Benjamini-Hochberg adjusted, linear-model derived P-value < 0.05 and fold-change > 1.25 or < 1/1.25). Grey circles indicate missing hits which were identified as significant in other detergent condition. Melting slopes for the entire dataset or significant PISA hits compared with Welch’s T-test. (H) Direct comparison of NP-40 and DDM PISA log2-fold changes: blue proteins are significant in DDM PISA alone, red proteins are significant in NP-40 PISA alone, and purple proteins are significant in both detergent PISA experiments. Dashed line represents equal fold changes for the two detergent PISA experiments.(J-K) PF-3758309-induced thermal shifts from NP-40 melt (J) or DDM melt (K) compared to thermal shifts from PBS melt^3^. Red points indicate significant PISA hit for NP-40 or DDM melt, respectively. Dashed line is where PBS PISA fold-changes equal detergent PISA fold-changes. (I) Selected proteins from D-E in barplot form for demonstrating cases where adding DDM to PISA melt buffer increases magnitude of PISA fold changes.

**Figure 4.**
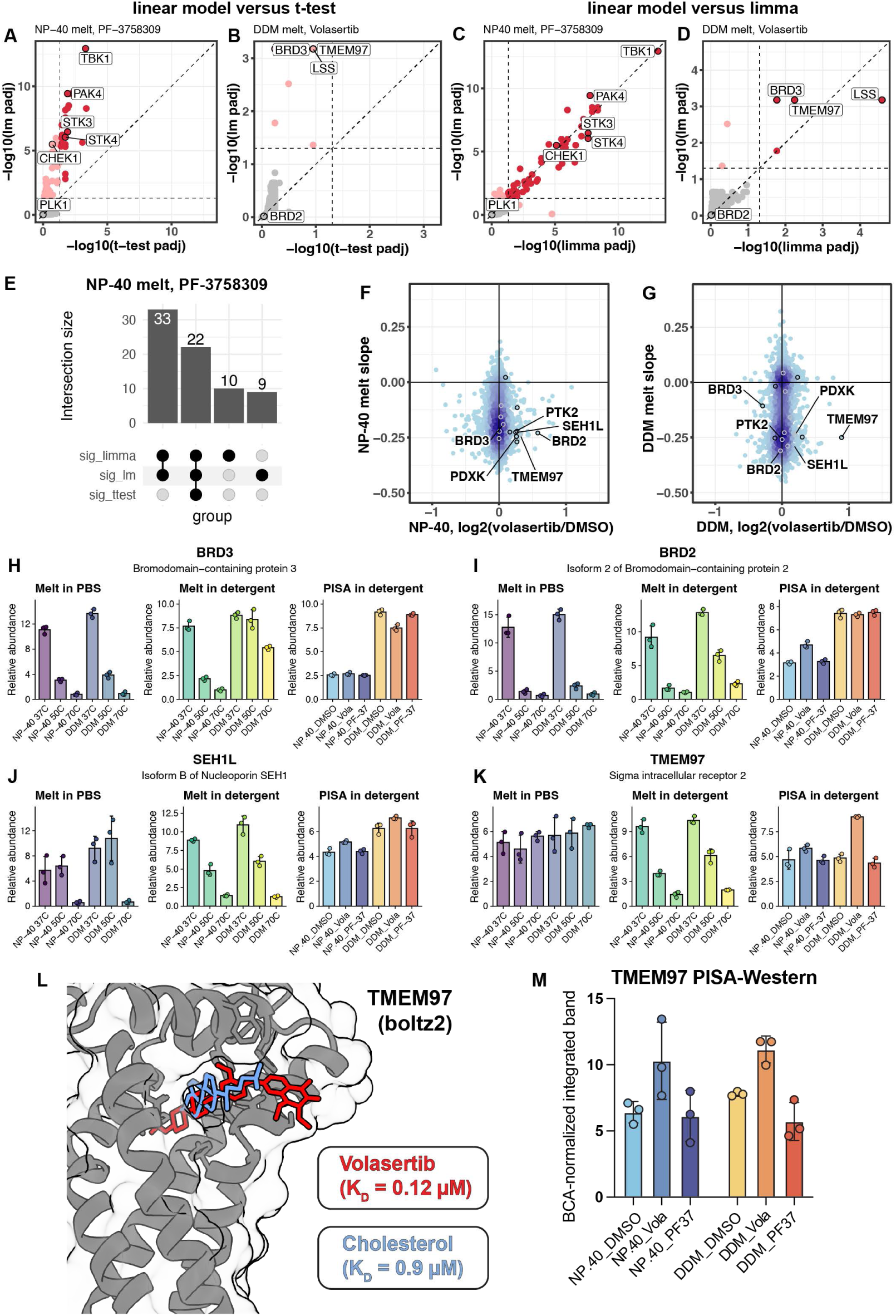
DDM identifies unique volasertib binding hits. (A-E) Comparisons of different methods for determining differential protein abundances. P-values are adjusted with Benjamini-Hochberge. Significance thresholds set at p-adjust < 0.05. Linear model approach (lm) compared to t-tests for PF-3758309 treatment in NP-40 detergent (A) or volasertib treatment in DDM detergent (B). Linear model approach compared to Limma-derived^24^ p-values for PF-3758309 treatment in NP-40 detergent (C) or volasertib treatment in DDM detergent (D). (E) Upset plot comparing significant hits for the Limma, linear model, and t-test methods. (F-G) Temperature B1 slopes from figure 2 plotted against volasertib-induced thermal shifts for NP-40 PISA (F) and DDM PISA (G). (H-K) Barplots comparing melting behavior in PBS, melting behavior in detergents, and PISA quantitative results in detergent for volasertib-destabilized BRD3 (H), volasertib-stabilized BRD2 (I), putative hits SEH1L (J) and TMEM97 (K). (L) Structure prediction in Boltz-2^34^ of TMEM97 with native ligand cholesterol (blue) or putative hit volasertib (red). (M) Densitometry integration of TMEM97 western blot of PISA samples (fig S7), normalized to total protein quantification by BCA.

To investigate how effectors of melting behavior affect PISA protein-ligand engagement detection, we compared the isothermal detergent temperature coefficient (β1) from our original experiment (**Fig. 2**) against significant “hits” in the PISA experiment (**Fig. 3F, G**). Proteins which were significantly altered in thermal stability with PF-37 treatment generally had steeper melting profiles in detergents, and PF-37 hits were generally confined to proteins that had measurable temperature-induced aggregation (β1 < 0) in the same detergent. Further, proteins which lost melting behavior in one detergent (β1 = 0) subsequently lost PISA hit identification. A clear example of this is in STK3: STK3 engaged PF-37 in the presence of NP-40, but measurement of this engagement was lost in the presence of DDM due to a loss of aggregation in the presence of DDM (**Fig. 3B**). In another case of melting behavior impacting PISA results, CHEK1 was destabilized when melted in the presence of both NP-40 and DDM, with stronger destabilization with PF-37 treatment and melting in the presence of DDM. DDM-dependent lowering of melting point resulted in a reduced initial abundance of CHEK1 (**Fig. 3C**). These examples highlight the phenomena that the position of the melting point relative to the temperature integration window alters our ability to measure a thermal shift in a positive (STK3) or negative (CHEK1) direction, in agreement with our model (**Fig. 1G**). While most PF-37 induced thermal shifts were identified in both NP-40 and DDM lysates, each detergent demonstrated unique hits, such as STK3, STK4, CHEK1, and TANK in NP-40, and MFSD5 in DDM (**Fig 3H**). Together, these observations underscore the importance of aggregation condition selection for PISA, and support a model that temperature-induced aggregation is a necessary, but not sufficient, condition for detecting drug–target engagement.

Through investigation of detergent effects on PISA engagement calling we observed that assays done in the presence of detergent, either NP-40 or DDM, increased PF-37-induced thermal shifts compared to performing the PISA in PBS followed by NP-40 extraction (**Fig. 3I-K**)^3^. In the case of PAK4, the annotated target of PF-37, PISA with DDM present during melting resulted in a 4-fold larger relative PF-37-induced stabilization of PAK4 compared to melting in PBS and 1.9-fold larger stabilization in DDM compared to NP-40 (**Fig 3I**). Beyond the annotated target of PAK4, CHEK1 is known to destabilize under PF-37 treatment^3^, but this destabilization was previously only significantly observed in an intact cell context. We find that performing the PISA in NP-40 recovers CHEK1 as engaging PF-37 in lysates, recapitulating previously observed in-cell destabilization behavior (**Fig 3I**). These observations of improved recovery of known PF-37 hits in lysate-PISA was encouraging but counterbalanced by the observation that some hits, like STK3 and STK4, were lost altogether due to the loss of temperature-induced aggregation in DDM (**Fig 3B**). The larger fold-changes in detergents in general could be due to several factors. NP-40 is known to broadly decrease melting points of proteins compared to melting in PBS alone^6^, an effect which is expected to increase PISA magnitudes (**Fig 1G**). Modeling a selection of PF-37 PISA hits in the Reinhard TPP dataset confirmed that in many cases NP-40’s melting point depression explains the difference in observed PISA fold-changes (**Fig S5**).

### Linear modeling approaches improve sensitivity of PISA assays

Thermal profiling assays such as PISA have found utility in screening-based experiments where the number of replicates is kept small to increase the number of conditions (e.g., drugs) tested, with larger screens using only duplicate measurements of drug-induced thermal shifts^3^. In addition, thermal shifts induce relatively small changes in protein solubility. Together, these effects make hit-calling in PISA assays particularly challenging and underpowered for traditional statistical testing. To overcome the limitations of t-tests, we investigated if multiple linear regression models combined with Benjamini-Hochberg correction to calculate adjusted p-values would improve hit recovery from PISA data. We grounded this idea based on the fact that pair-wise t-tests only use the given two comparisons to estimate variance, while a linear model across multiple conditions uses one model for each protein, which enables the use of all samples within an experiment to estimate variance for that protein^31^. This approach leverages an important advantage of TMT-based proteomics which is the simultaneous preparation and quantitation of all samples in a plex. Comparing linear-model p-value calculations to t-test p-value calculations, we found that linear models outperformed t-tests across different experimental conditions (**Fig. 4A-B**). In addition, we compared simple linear modelling with the well-established Limma method, which borrows variances across genes for differential gene expression estimation^24^. In both drug-treatment assays, Limma and the simple linear model approach achieved similar performances, with either linear model technique demonstrating improved performance over t-tests. (**Fig 4C-E**). This similarity demonstrates the viability of a simple linear model approach to differential abundance tests. Encouragingly, both methods recover BRD3 as a significant volasertib hit in DDM lysate, and additionally identify a previously unreported interaction with the cholesterol-binding membrane protein transmembrane protein 97 (TMEM97; sigma-2 receptor) (**Fig 4D**). To assess the sensitivity of linear modeling in the context of high-throughput screens with fewer replicates, we compared linear model p-values in a PISA experiment titrating the PLK1 inhibitor BI2536 from 100nM to 10μM using either 4 DMSO controls (**Fig. S6A-D**) or 2 DMSO controls (**Fig. S6E-H**). Encouragingly, multiple linear regression consistently recovered PLK1 as a PISA hit even at 100nM, the lowest concentration tested. Consistent with the improved statistical power stemming from pooled variance estimates, PLK1 was identified as a hit, even when the number of DMSO controls was down sampled to 2 replicates. These findings demonstrate that a linear modeling, either with canonical linear models or limma, can enhance the sensitivity of PISA assays for detection of known drug targets even at concentrations much lower than those used previously in high-throughput PISA assays^3^.

### DDM and NP-40 identify unique PISA hits upon treatment with volasertib

The PLK inhibitor volasertib is known to also inhibit bromodomain proteins BRD2 and BRD3. However, these interactions were previously only observed when PISA was done on intact cells, not on cell lysate^3^. We find that DDM preferentially stabilizes the melt behavior of BRD2 and BRD3, with consequences for PISA hit calling. DDM stabilization allowed for the detection of a volasertib-induced destabilization of BRD3 (**Fig 4H**), but masked a volasertib-induced stabilization of BRD2 (**Fig 4I**). However, both DDM and NP-40 facilitated similar improvements to the melting behavior of the nuclear pore protein SEH1L, consistent with our findings that both detergents improved the melting behavior of membrane proteins (**Fig 2G-H**) and that DDM particularly improved nuclear pore protein melting behavior (**Fig 2I**). Subsequently, volasertib induced a similar stabilization of SEH1L in both detergents (**Fig 4J**) consistent with previous findings that SEH1L interacts with BRD2 in AP-MS experiments^32^. Additionally, volasertib induced significant thermal shifts in NP-40 lysate for the kinases PDXK and PTK2 (**Fig 4F**), which have been previously unidentified in PBS lysates but are known volasertib hits in a kinobead assay^33^.

Isothermal protein melts indicated that membrane proteins exhibited more temperature-induced aggregation when melted in detergent, (**Fig 2G-H**) which we demonstrate can improve PISA hit detection. Consistent with this, TMEM97 was identified as a volasertib hit only in the presence of DDM (**Fig. 4D, F-G**), and TMEM97 itself exhibited temperature-induced aggregation only when detergent was present during the melt (**Fig. 4K**). Computational modeling of the volasertib–TMEM97 interaction with Boltz-2^34^ confirmed TMEM97 binding of its endogenous ligand cholesterol (predicted affinity: cholesterol: IC50 = 0.9 µM). In addition, Boltz-2 predictions of TMEM97 with volasertib predicted that volasertib could occupy the same cholesterol binding pocket but with a lower predicted affinity (predicted affinity: volasertib: IC50 = 0.12 µM). We subsequently validated this engagement based on PISA experiments with orthogonal Western blot readout (**Fig. 4L-M, Fig. S7**). Overall, the TMEM97 finding supports our model of proper melting behavior, as defined by a steep melting slope (**Fig. 1E**) and near-zero minimum value (**Fig. 1D**), are required for the identification of a drug-target hit in this protein.

## Conclusions

Lysate-based and intact-cell PISA experiments return different sets of hits, with intact-cell assays typically yielding more^3^. This gap is generally attributed to the preservation of native cellular context—downstream effectors of drug binding, endogenous cofactors such as ATP^1^, and could be due to the stabilization effects of intact membrane environments^6^. Our findings suggest that effects usually ascribed to “cellular context” can be reconstituted, at least partially, by judicious choice of assay conditions, including detergents. The rescue of TMEM97 as a volasertib hit in DDM but not NP-40 demonstrates this concept, wherein a transmembrane protein that fails to aggregate meaningfully in one detergent becomes tractable in another, enabling a ligand-induced shift to be measured.

While our study points to key considerations for protein-ligand engagement detection with thermal stability assays, our study has several limitations. We examined two detergents at a single concentration in one cell line. Broader panels of detergents, concentrations, and cell backgrounds will be needed to generalize these conclusions. Our hit-calling analysis was limited to two compounds selected precisely because of their known in-cell versus lysate behavior. A wider compound survey will be required to quantify how often detergent choice is decisive for PISA outcomes.

These findings carry two practical implications for thermal proteome profiling. First, detergent choice should be treated as an experimental variable rather than a fixed methodological detail. Reports of drug–target engagement, or the absence thereof, for membrane-associated targets in lysate-based formats warrant re-evaluation under alternative detergent conditions before strong negative conclusions are drawn. Finally, these results argue for deliberate buffer optimization when designing PISA (or related TPP experiments), particularly for compounds expected to engage membrane or membrane-associated proteins.

## Acknowledgements

The authors would like to thank the Schweppe Lab for their helpful advice and comments. This work was supported by: R35GM150919 (DKS), the W.M. Keck Foundation (DKS), an Andy Hill CARE Distinguished Researcher Award (DKS), a Cancer Consortium New Investigator Award (funded in part through P30 CA015704, DKS), and the Pew Charitable Trusts (DKS).

## Conflicts of interest

The authors declare the following competing financial interest(s): D.K.S. is a consultant and/or collaborator with ThermoFisher Scientific, AI Proteins, and Genentech. The other authors have no conflicts to declare.

## Supplementary Information

**Supplemental Figure 1.** Four-parameter fit of melting curves identifies outlier melting behavior.

**Supplemental Figure 2.** Relative qualitative behaviors of various detergents in melting buffers for thermal profiling.

**Supplemental Figure 3.** Different metrics to evaluate melting profiles of proteins from isothermal melts.

**Supplemental Figure 4.** Melting profile bar graphs for selected proteins from pfam and Jensen’s compartments terms.

**Supplemental Figure 5.** Four-parameter melting curve fit to Reinhard, 2015 datasets for TPP in PBS or 0.4% NP-40.

**Supplemental Figure 7.** Western blot validation of TMEM97.

## Notes

https://github.com/SchweppeLab/DDM_PISA_paper

### Summary of Updates

In our revised manuscript we (a) replicated our PISA findings with a dataset that no longer contained outlier samples, (b) added figure 3H directly comparing PISA results of the two detergents, (c) fixed typos in manuscript and supplemental information, and (d) altered title from "PISA" to "thermal stability proteomics" to improve accessibility.

