## Supplemental figures 1-7 for "Detergent selection as a determinant of protein melting behavior and drug-target interaction-calling in thermal stability proteomics"

### Determinants of lysate melting behavior and drug-target hit-calling in thermal stability proteomics

#### *Supplementary Figures*

**Supplemental Figure 1.** Four-parameter fit of melting curves identifies outlier melting behavior.
**Supplemental Figure 2.** Relative qualitative behaviors of various detergents in melting buffers for thermal profiling.
**Supplemental Figure 3.** Different metrics to evaluate melting profiles of proteins from isothermal melts.
**Supplemental Figure 4.** Melting profile bar graphs for selected proteins from pfam and Jensen's compartments terms.
**Supplemental Figure 5.** Four-parameter melting curve fit to Reinhard, 2015 datasets for TPP in PBS or 0.4% NP-40.
**Supplemental Figure 6.** Linear modeling outperforms t-tests in low-N screening experiments.
**Supplemental Figure 7.** Western blot validation of TMEM97.

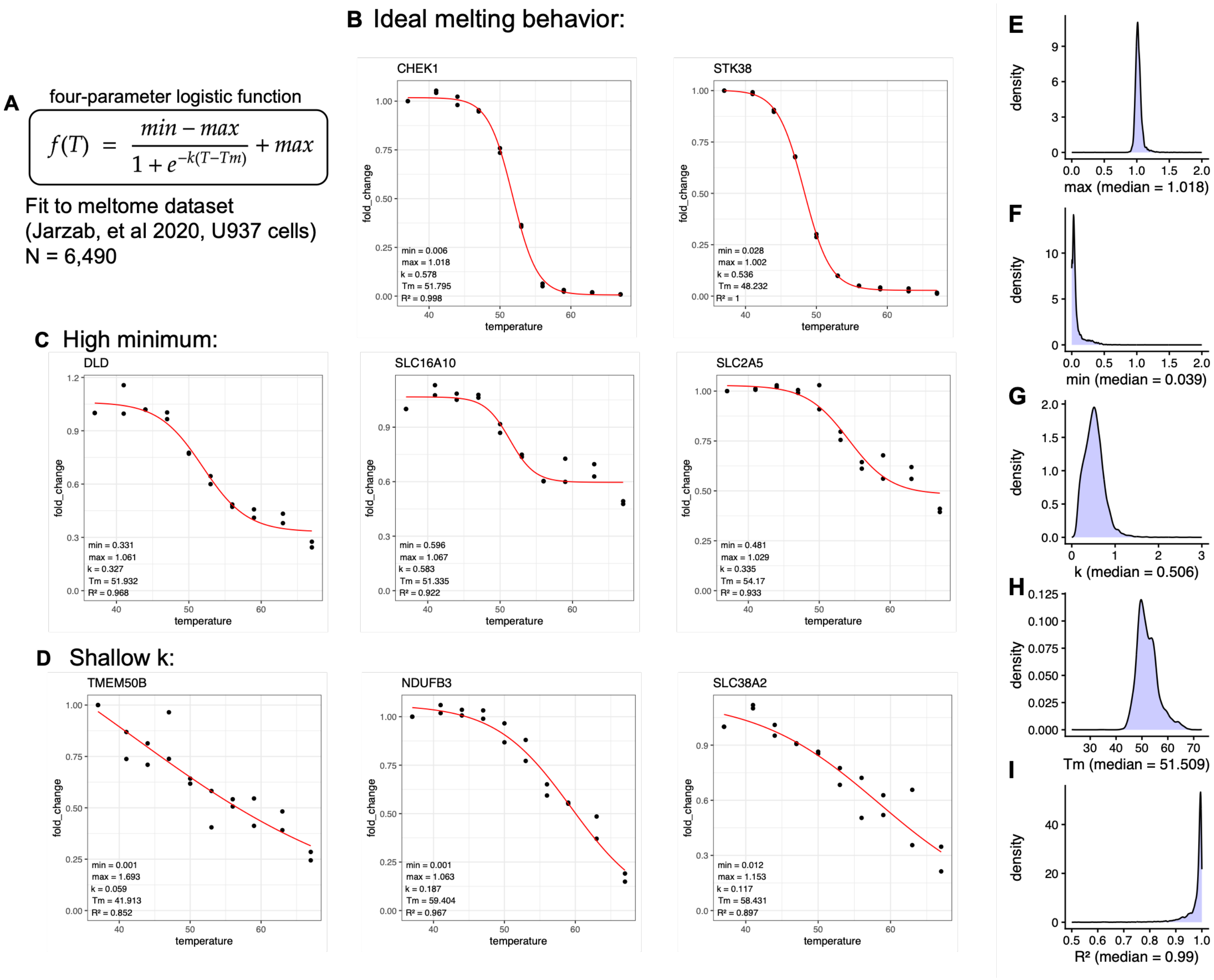


**Supplemental Figure 1. Four-parameter fit of melting curves identifies outlier melting behavior.** (A) Formula defining protein melting behavior as a function of minimum signal, maximum signal, melting point Tm and slope constant k. (B) Examples of well-behaved proteins with good curve fit, minimum signal around 0, maximum signal around 1, and steep transition (high k). (C) Examples of proteins with high minimum signal. (D) Examples of proteins with shallow transitions (low k).

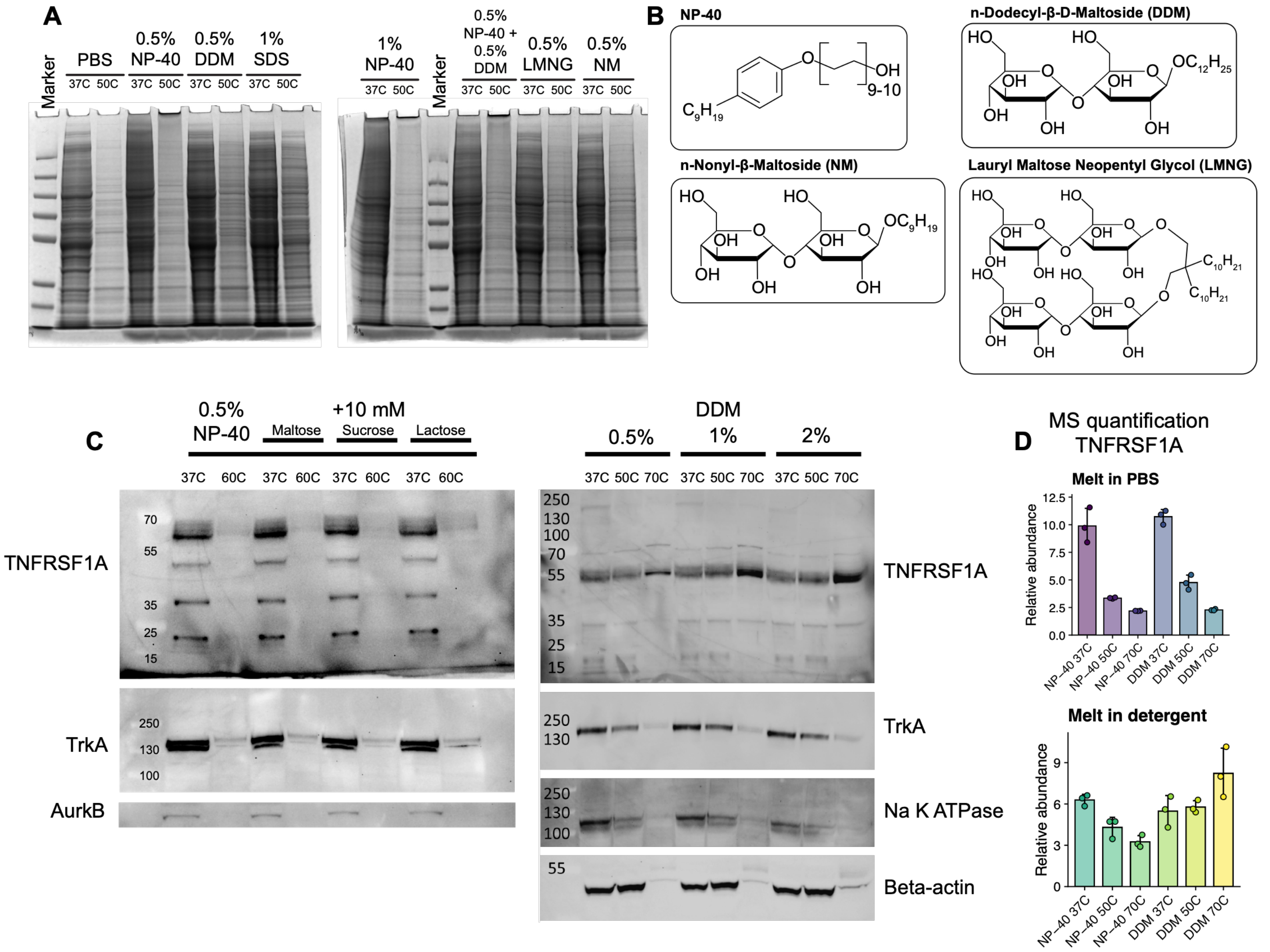


**Supplemental Figure 2. Relative qualitative behaviors of various detergents in melting buffers for thermal profiling.** (A) SDS-PAGE gel of soluble fraction from TF-1 cell lysate prepared with indicated detergent additive and heated to indicated temperature. (B) Chemical structures of NP-40 and maltoside detergents used in this work. (C) TF-1 cell lysates were prepared with indicated additive in PBS and heated to indicated temperature. Lysates were clarified by centrifugation and clarified lysate analyzed by western blot for various proteins of interest. (D) Mass spectrometry quantification of TNFR1 confirming melting behavior impacts of DDM.

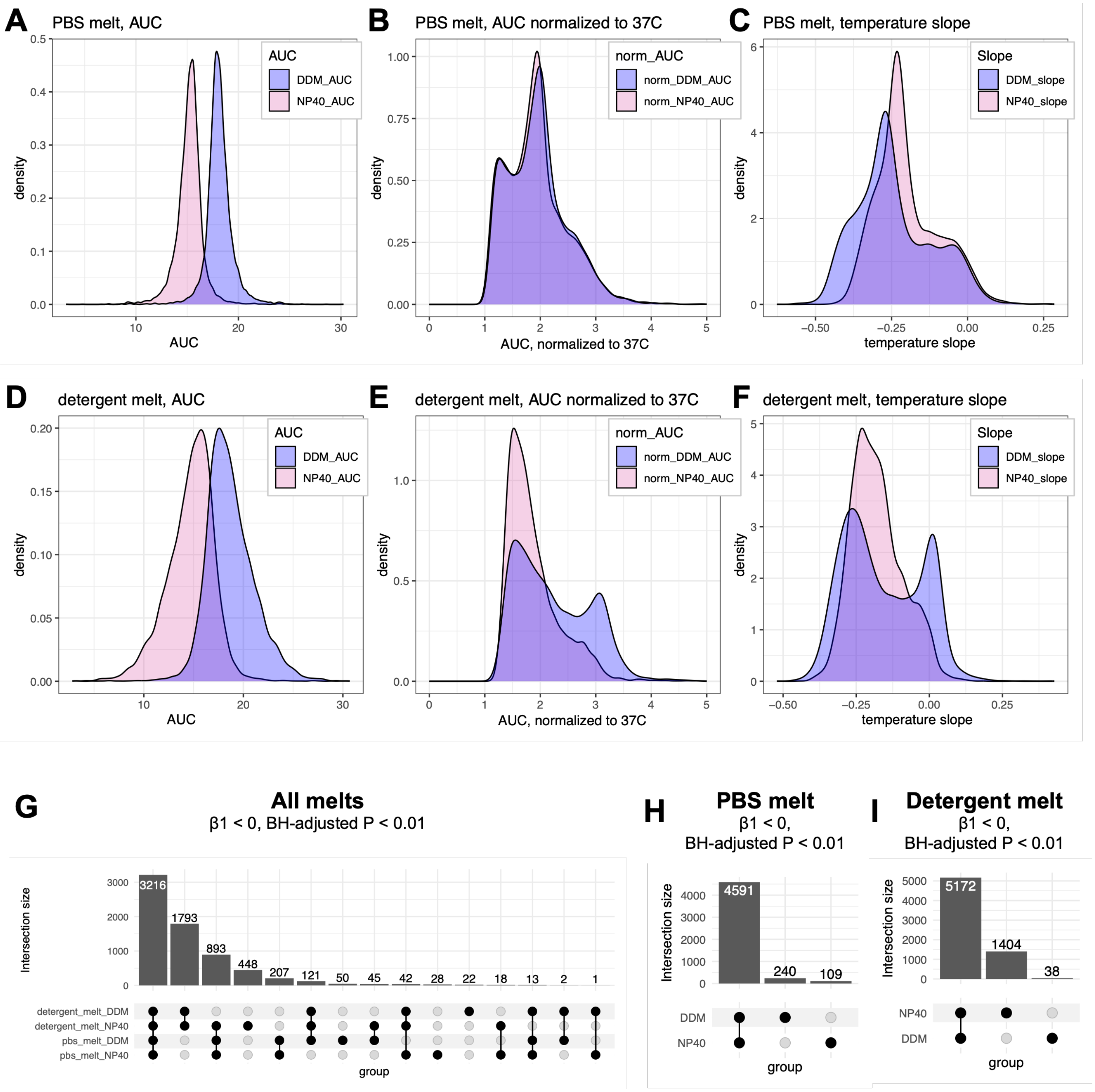


**Supplemental Figure 3. Different metrics to evaluate melting profiles of proteins from isothermal melts.** (A, D) Area under the curve (AUC) metric for PBS melt (A) and detergent melt (D) obtained by summing each temperature point for each protein and detergent condition. (B, E) AUC normalized to 37 degree temperature point for each protein from PBS melt (B) and detergent melt (E). (C, F) Temperature slopes of each protein calculated from linear regression for PBS melt (C) and detergent melt (F). (G-I) Counts of significant slopes (linear regression, Benjamini-Hochberg adjusted p-value < 0.01, temperature slope β1 < 0) across all melt conditions (G), PBS melt followed by detergent extraction alone (H) or detergent melt alone (I).

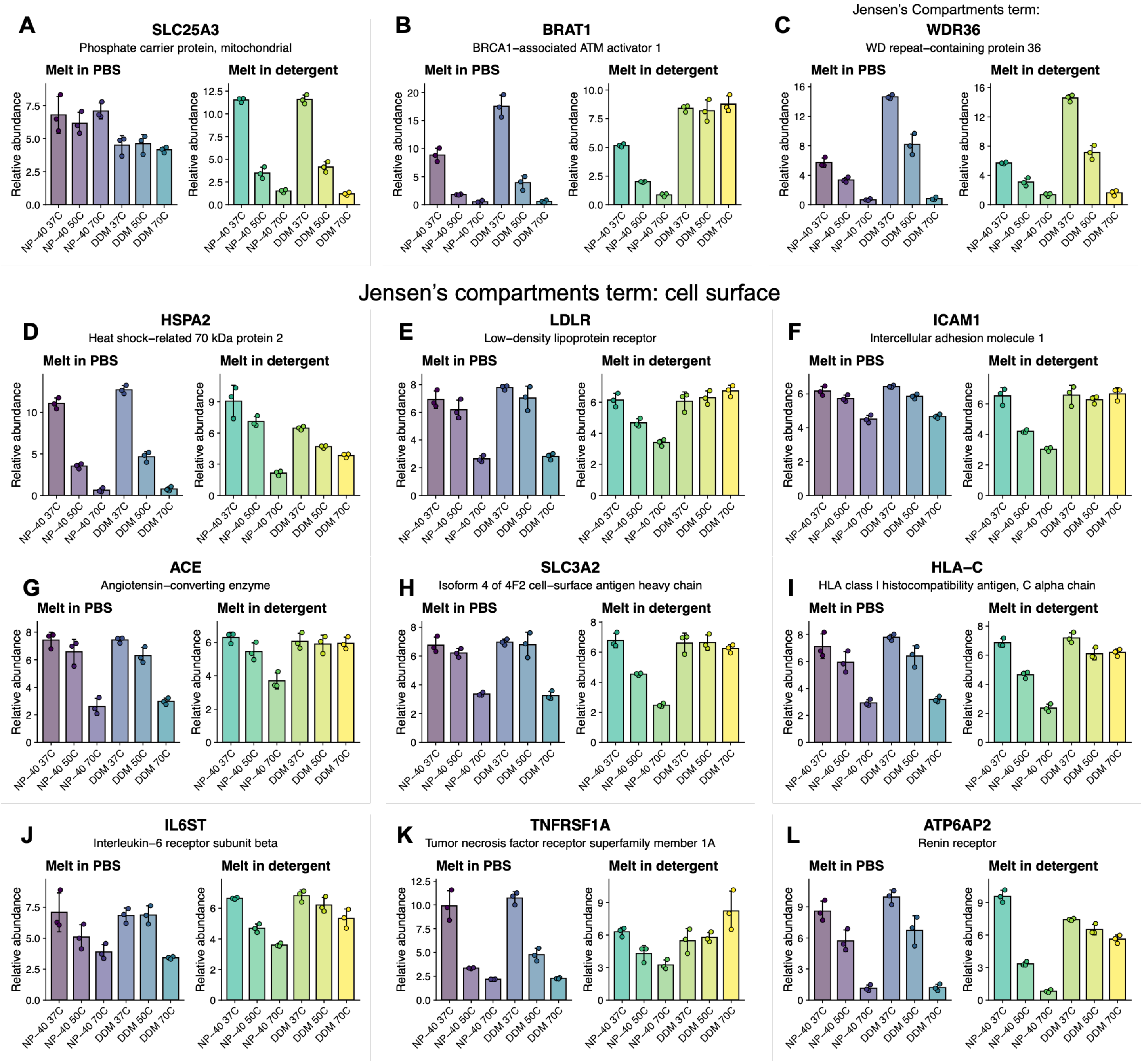


**Supplemental Figure 4. Melting profile bar graphs for selected proteins from pfam and Jensen's compartments terms.** (A) SLC25A3 which melts only in the presence of detergent. (B) BRAT1, which melts without detergent and in the presence of NP-40, but not DDM. (C) WDR35 which has similar melting profiles with or without detergent, but higher abundance in DDM. (D-L) Selection of cell surface proteins which have little to no melting behavior in DDM compared to NP-40.

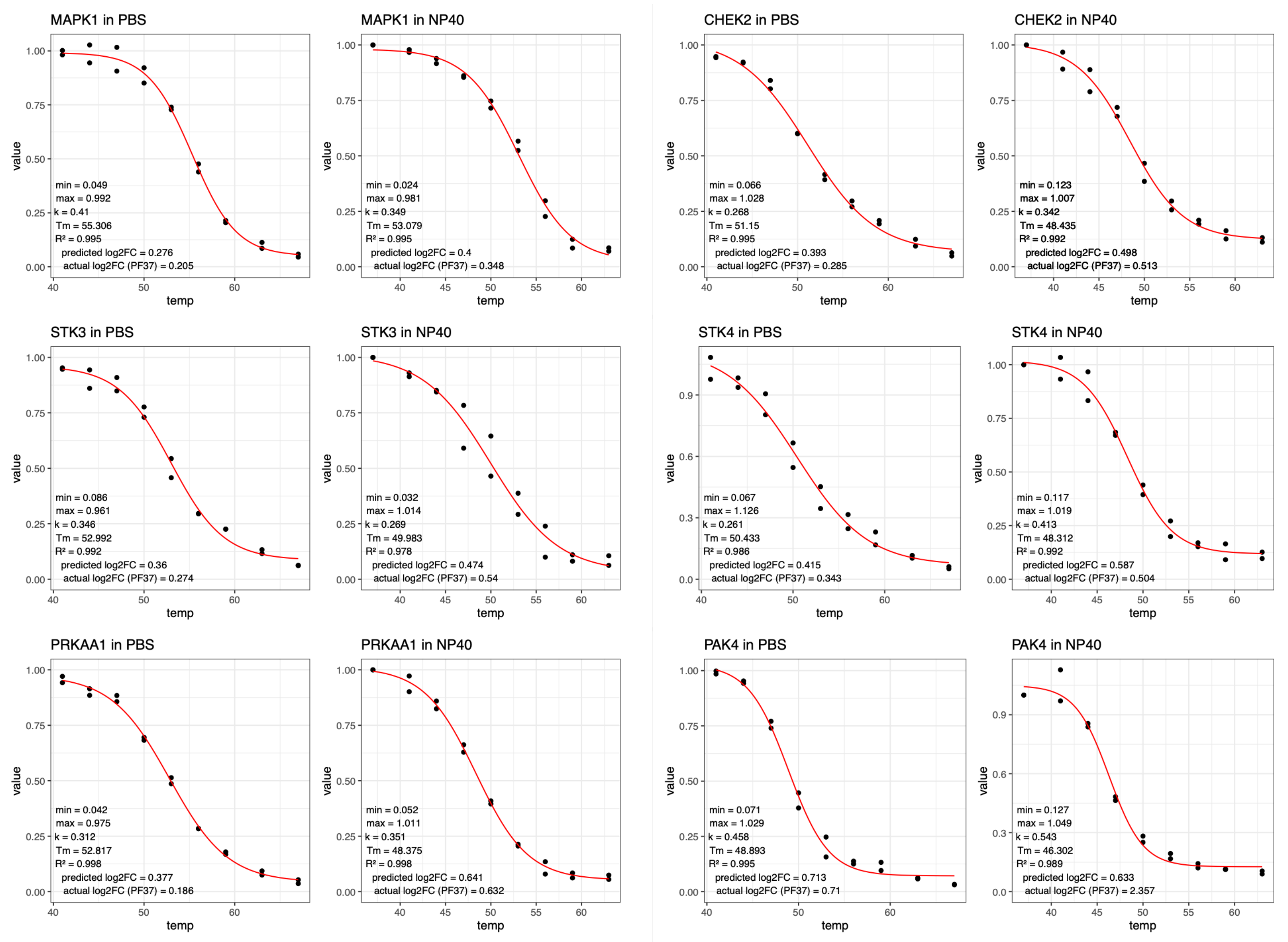


**Supplemental Figure 5. Four-parameter melting curve fit to Reinhard, 2015 datasets for TPP in PBS or 0.4% NP-40.** Selected proteins from PF-3758309 PISA hits. Predicted log2FC calculated by integrating area under fit curve and area under fit curve with +2.5C added to Tm and taking log2FC(Tm+2.5/Tm). Actual log2FC values taken from PBS melt dataset(Vranken et al. 2024) or NP-40 melt dataset (this work).

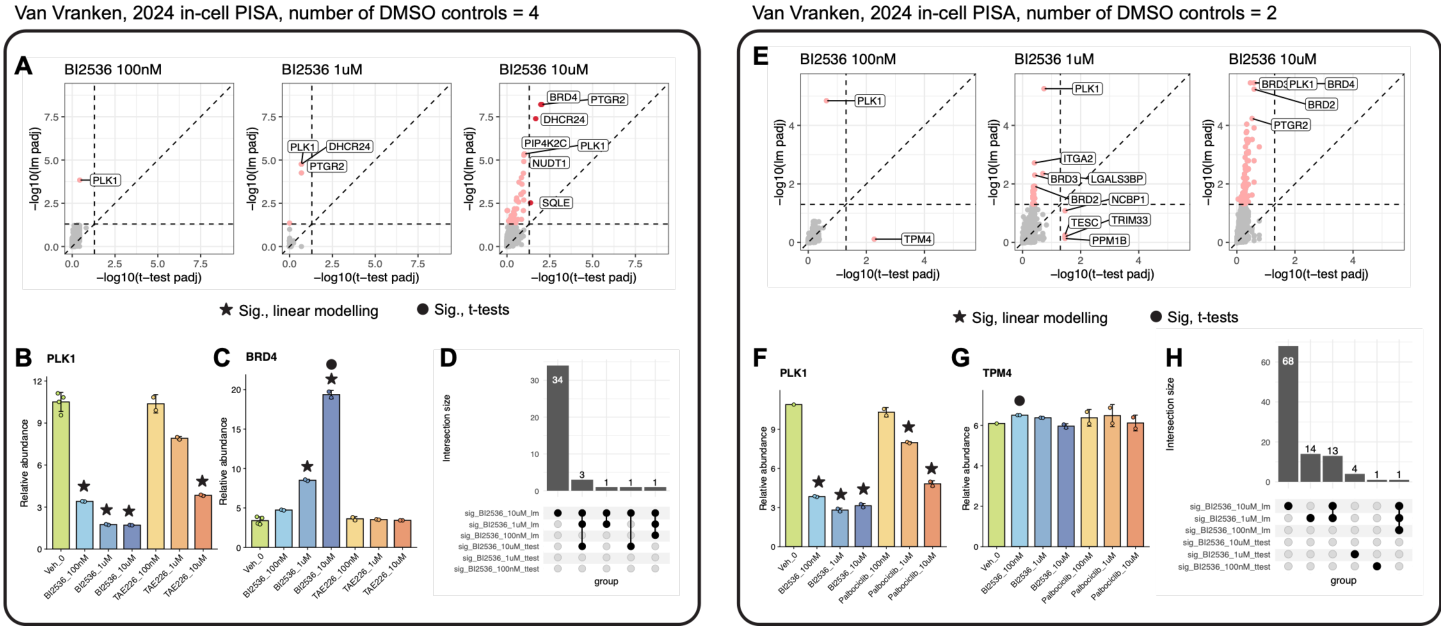


**Supplemental Figure 6. Linear modeling outperforms t-tests in low-N screening experiments.** Dose-response PISA from Van Vranken 2024 used to benchmark linear modeling approach. (A, E) Benjamini-Hochberg adjusted, -log10(p-value) calculated by linear model approach or Welch's t-test for two independent experiments, one with treatment N = 2 and control N = 4 (A) and another with treatment N = 2 and control N = 2 (E). (B-C, F) selected proteins demonstrating sensitivity of lm approach. (D, H) counts of significant proteins for the three concentrations and two approaches. (G) T-test unique hit shown to demonstrate t-test sensitivity to low-N artifacts.

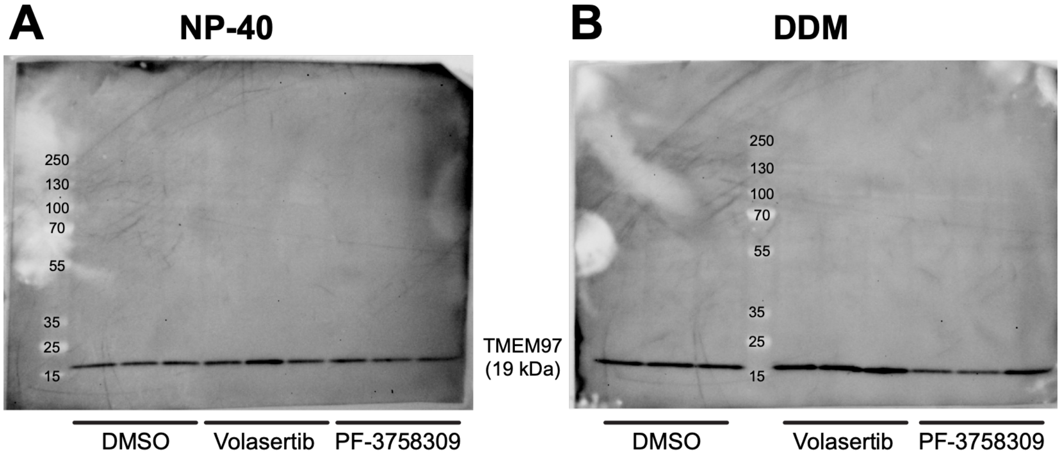


**Supplemental Figure 7. Western blot validation of TMEM97.** Clarified lysate from detergent PISA blotted with anti-TMEM97 antibody (CST 62790T) and visualized with anti-rabbit IgG HRP-linked antibody (CST 7074P2).
